# The Role of Reproductive Periodicity in Dispersal Among Hydrothermal Vents and its Implications for Regional Connectivity and Conservation

**DOI:** 10.1101/2023.03.07.531641

**Authors:** Otis Brunner, Pierre Methou, Satoshi Mitarai

## Abstract

Connectivity among isolated habitat patches via planktonic larval dispersal is crucial for maintaining the regional diversity of hydrothermal vents. Despite, increasing sophistication of techniques for simulating dispersal, limited information on biological and behavioural traits of vent-associated species has unknown affects on the applicability of these methods for conservation. Here we focus on the role of periodic reproduction on dispersal among hydrothermal vents, as periodic spawning has increasingly been observed in a variety of taxa. For generalizeability, we simulate the dispersal of larvae under treatments of periodic and aperiodic release timing at various depths, with a consistant but variable planktonic larval duration. Our results show a highly variable effect of periodicity on the characteristics and distribution of dispersal, which are heavily modified by the dispersal depth and source location. The capacity for reproductive periodicity to impact the among-site dispersal warrents further investigation into its prevelance and timing among vent-associated fauna.

## Introduction

Connectivity is the process by which populations and communities at distant habitat patches interact through the sharing of individuals, to the extent that these isolated populations/communities influence each other’s demographic processes (Crooks and Sanjayan 2006; Correa Ayram et al. 2016). Connectivity at this scale supports the local and regional diversity of the habitat in question as well as providing resilience to disturbance events. For these reasons, connectivity and its maintenance are considered central to conservation objectives (IUCN, 2017). With a high sensitivity to natural disturbance, due to their ephemeral nature and high degree of isolation, connectivity in hydrothermal vents, is an essential aspect of a species’ survival over successive generations (Mullineaux *et al*., 2018). Added to that, are the growing anthropogenic pressures at these ecosystems, with the prospect of deep-sea mining in the foreseeable future (Van Dover 2014; Turner *et al*., 2019).

Connectivity among hydrothermal vents is difficult to quantify or predict because it requires knowledge on several sequential steps in a species’ life history, from reproduction at the source site to successful colonization at the target site, all of which are influenced by a complex interaction of biotic and abiotic factors (Kritzer and Sale 2004; Pineda et al. 2007; Thompson *et al*., 2020). Genetic models using multiple markers are often used as evidence for direct or indirect connectivity among distant hydrothermal vent sites based on the ongoing or historical transfer of genetic material (reviewed by Vrijenhoek, 2010). However, the process of dispersal, which facilitates genetic connectivity, is inherently difficult to observe directly. Thus, larval transport among hydrothermal vents has often been predicted using hindcast velocity fields within models of ocean circulation (Mitarai *et* al., 2016; Breusing *et al*., 2016; Breusing *et al*., 2021). To simulate dispersal within these ocean circulation models, it is necessary to incorporate the biological and behavioral traits of larvae, such as the planktonic larval duration (PLD), vertical position, spawning behavior etc. (Swearer *et al*., 2019; Jahnke and Johnson, 2021). There is considerable uncertainty surrounding dispersal behaviour of deep-sea species (reviewed by Hilario *et al*. (2015)), but generalizations of dispersal behavior have been used to answer important questions using a range of dispersal abilities supported by the best available biological and oceanographic data (Young *et al*., 2012; Mitarai *et al*., 2016; McVeigh *et al*., 2017; Breusing *et al*., 2021). As in other marine ecosystems (Cowen and Sponaugle, 2009; Treml *et* al., 2015), these studies have highlighted the importance of the PLD and vertical position at hydrothermal vent ecosystems, by testing a range of different dispersal depths and larval duration. Although it is common for simulations of dispersal to ignore temperature dependent growth rates, instead using a fixed PLD (Swearer *et al*., 2019), previous works have contradicted these assumptions for vent ecosystems and showed that ocean currents and temperature have an interacting effect on dispersal at different depths (Mitarai et al., 2016; Breusing et al., 2021). In addition, Yahagi *et* al. (2017) also demonstrated the importance of temperature dependent growth in *Shinkailepas myojinensis* larvae dispersing from hydrothermal vents in the Northwest Pacific.

Conversely, the influence on dispersal of other life-history traits, such as those related to reproductive strategies, have been poorly investigated in hydrothermal vent species. Yet, both timing and frequency of spawning have been shown to drastically impact connectivity of species from coral reefs and coastal ecosystems (Carson et al. 2010, Kough & Paris 2015, Romero-Torres et al. 2017); (reviewed by Swearer *et al*., 2019)). Moreover, surface generated mesoscale eddies potentially result in the seasonal and interannual variability of larval transport from hydrothermal vents (Adams et al., 2011), which could be exacerbated by variations in spawning behavior. For example, species that reproduce synchronously at a certain time in the year (periodically) may be more strongly influenced by annual and inter-annual variability than those that reproduce throughout the year (aperiodically).

Endemic fauna from hydrothermal vents exhibit a wide range of reproductive strategies and spawning behaviors, similarly to species from other marine ecosystems. Although most have aperiodic reproductive patterns — sometimes defined as continuous — (Jollivet et al., 2000; Faure et al., 2007; Tyler et al., 2008; Hilário et al., 2009; Matabos and Thiebaut, 2010; Nakamura et al., 2014; Marticorena et al., 2020), cases of periodic reproduction have also been observed in some families despite the absence of sunlight at these depths (Perovich et al., 2003; Dixon et al., 2006; Methou et al., 2022). In *Bathymodiolus* mussels from the Atlantic and some bythograeids crabs from the East Pacific, reproductive periodicity correlates with seasonal variations of the photosynthetic primary production sinking from the surface, with a spawning or hatching period preceding the peak of photosynthetic production (Perovich et al., 2003; Dixon et al., 2006; Tyler et al., 2007). Unlike all these previous works, the intriguing case of *Rimicaris exoculata* and its sister species *R. kairei* revealed a brooding phase restricted to the period between January and early April, regardless of the hemisphere and therefore without any apparent relationship with a known seasonality pattern (Methou et al., 2022). This contrasts with reproductive rhythms in other shrimps from the same family, with brooding phases that match seasonal variations from the surface (Copley and Young, 2006) or are simply aperiodic (Methou et al. in review). Thus, periodic and aperiodic reproduction sometimes coexist in species of the same family (Perovich et al., 2003; Copley and Young, 2006; Hilário et al., 2009; Methou et al., 2022), resulting in different timing of larval release among conspecifics.

In this paper we investigate the role of reproductive periodicity on dispersal by comparing simulations of year-round spawning to simulations where spawning is restricted to a single month. Although it has been found that the stochasticity of surface currents means that periodicity will have little effect on connectivity in coastal ecosystems on the annual scale (Siegel *et al*., 2008) this has not been tested for deep-sea ecosystems. It is not unreasonable to assume that dispersal in deeper waters should result in less stochasticity and therefore a more consistent inter-annual signal of reproductive periodicity.

The geographical extent of these simulations is limited to an area of the Northwest Pacific which represents a distinct bioregion (Bachraty et al. 2009) with little influence from dispersal from external sites (Mitarai *et al*., 2016). Local and regional diversity is particularly high in the Northwest Pacific compared to other hydrothermal vent regions (Thaler and Amon, 2019) and species distributions are somewhat spatially structured (Brunner *et al*., 2022) within the region. This region is also home to the world’s first full-scale test mining of a hydrothermal vent ecosystem, which was carried out in the central Okinawa Trough in 2017 (Okamoto *et al*., 2019). The regional impacts of localized disturbances has been poorly studied in the Western Pacific (Mullineaux *et al*., 2018) and the only such study to attempt this so far (Suzuki *et al*., 2018) utilized dispersal estimates that assumed aperiodic dispersal (Mitarai *et al*., 2016) and do not account for any variability associated with larval release timing. Currently, periodic reproduction has not yet been clearly reported in vent species from the Northwest Pacific, however this region hosts several vent species – including bathymodiolin mussels, alvinocaridid shrimps or bythograeid crabs –, with known cases of seasonality in congeners from others bioregions (Perovich et al., 2003; Dixon et al., 2006; Methou et al., 2022). Aside from release timing, we kept the important parameters of dispersal depth and PLD in our simulations mostly consistent with those used in previous studies (Young *et al*., 2012; Mitarai *et al*., 2016; Breusing *et al*., 2021). In this study we test whether or not reproductive periodicity influences the dispersal ability of species from hydrothermal vents and how this interact with other dispersal parameters such as depth of dispersal. Furthermore, we investigate how periodicity and depth affect the applicability of certain metrics used to assess regional connectivity and inform conservation.

## Methods

### Study Sites

The hydrothermal vent sites that are the focus of this study are mostly distributed within the Okinawa Trough, Izu-Bonin Arc, Mariana Arc, and Mariana Trough (Figure 1). We considered sites listed in InterRidge ver. 3.4 (Beaulieu and Szafranski, 2020) as active and confirmed as viable ‘source sites’ for the dispersal of larvae. We added several hydrothermal vent sites that are not included in InterRidge ver. 3.4 but have been recently confirmed (Methou et al. in review) as well as a number of cold-seep ecosystems that have been found to share species with hydrothermal vents in the region (Tokuda *et* al., 2006; Feng *et al*., 2018; Xu *et al*., 2018). A list of the sites with associated metadata can be found in Supplemental Table 1 and a map of their locations in Supplemental Figure 1. The dominant oceanographic features within the extent of this study are the Kuroshio western boundary current and the North Equatorial Current (NEC) (Figure 1).

**Figure 1.**
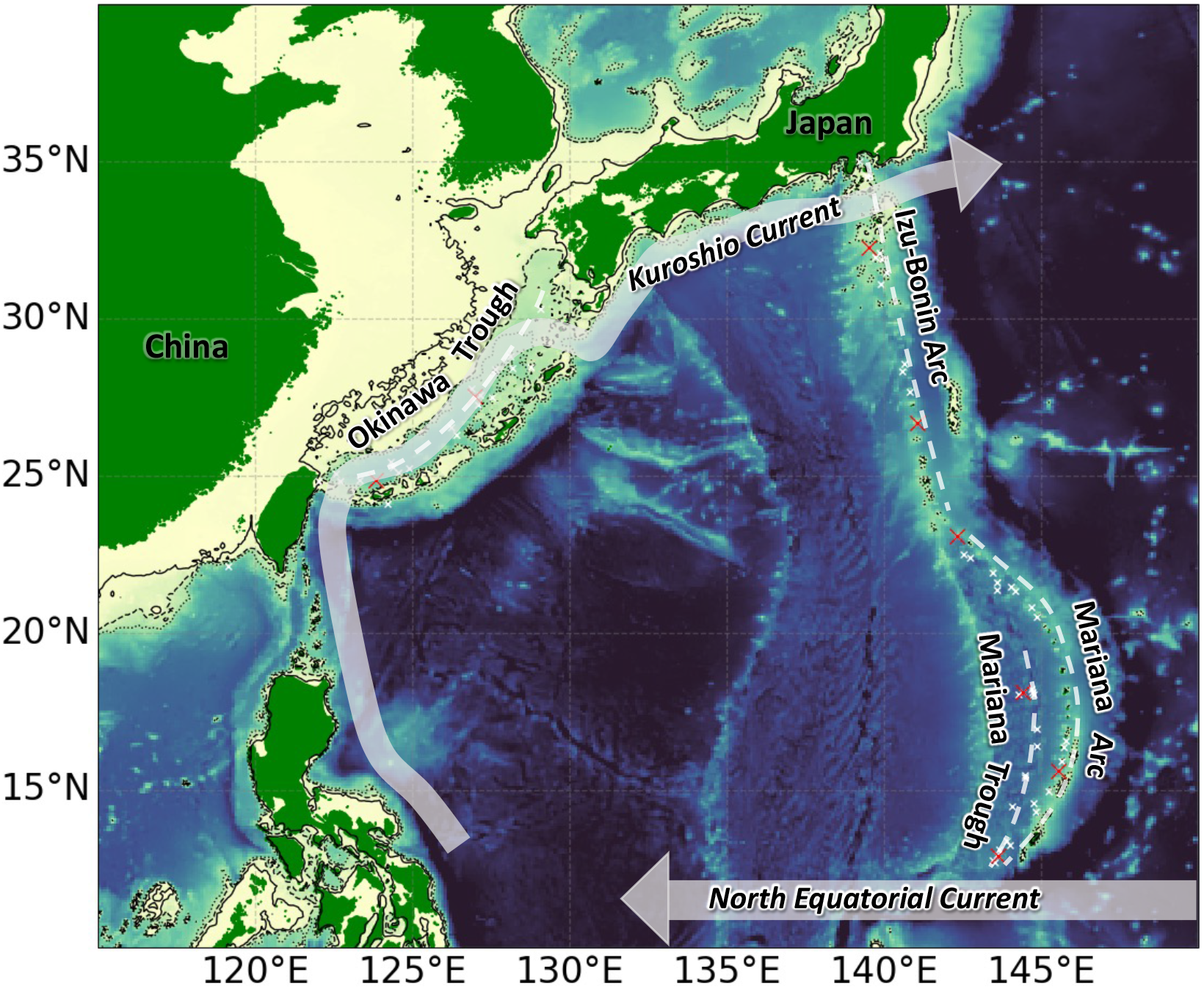
Study site in the Northwest Pacific showing the distribution of vent sites (red ‘x’ for focal sites and white for the rest) on their respective tectonic features (dashed white lines) and the approximate location of the two most dominant currents in the region (grey arrows). The colourmap is based on bathymetry data from ‘MOI GLORYS12_FREE’ 1/12^th^ ° Ocean General Crculation Model with the contours plotted as black lines at 100m (solid line), 500m (dashed line), and 1000m (dotted line).

### Dispersal Simulations

All simulations of larval dispersal, as well as the analyses and plotting of outputs, were carried out using Python version 3.10.2 (http://www.python.org) and PARCELS version 2.3.1 (Delandmeter and Sebillie, 2019). PARCELS is a set of methods for lagrangian particle tracking, which use oceanographic velocity data as an input, and track the advection of simulated particles based on this velocity data and subgrid-scale diffusion (Smagorinsky, 1963). The velocity data was obtained from the ‘MOI GLORYS12_FREE’ Ocean General Circulation Model via MERCATOR. Temperature data was also obtained from the same source. The resolution of the velocity and temperature field data used was 1/12^th^ of a degree horizontally with daily timesteps across 50 depth bands from 0.5m to 5728 m depth. The temporal extent of the data was five years from 1997 to 2003 in order to the minimize the affect of Kuroshio ‘large meandering events’ (Qui and Chen, 2021) on inter-annual variability. The spatial extent of the data covered the entire North Pacific Ocean to eliminate the boundary effects of larvae dispersing to the edge of this extent.

Using the same oceanographic field data, we ran simulation under three hypothesized dispersal depths; 100m, 500m, and, 1000m. In each simulation, lagrangian particles (with behaviors that approximate planktonic larvae) were released simultaneously from the same 69 source sites around the Northwest Pacific. In cases where the maximum depth of the oceanographic data at a source site exceeded the depth of dispersal, the output from that source site was not included in analyses. This resulted in a total of 69 sites for 100m, 66 sites for 500m, and 63 sites for 1000m scenario (Supplemental Table 1). 20 lagrangian particles were simultaneously released from each source site every six hours during the release window dictated by the periodicity of larval release. The total number of particles released from each site was 146,000 across the five years and roughly 2433 (24,6000/60) for each month release within the five years. The number of days from release to termination/settlement (PLD) for each larva was individually calculated based on the seawater temperature they encountered at hourly time intervals. The equation to calculate PLD from temperature was adapted from O’Connor *et al*. (2007) following the methods of Mitarai *et al*. (2016) and Breusing *et al*. (2021). Based on O’Connor *et al*. (2007) finding that the magnitude of the PLD-temperature relationship varied for each of the 69 species they tested, we used the highest magnitude which they found to suitably fit this relationship. We therefore assume that deep-sea hydrothermal vent species have a relatively long PLD compared to the shallow-water species tested by O’Connor *et al*., (2007). For reproducibility and consistency with other studies, we followed the methods recommended by the creators of PARCELS as closely as possible. The details of the simulation parameters along with links to scripts and tutorials can be found in Supplemental material.

### Analysis

In the simulations we recorded each lagrangian particles’s age, average temperature experienced, and coordinate position at the end of its PLD. We analysed the variability of these outputs in response to the addition of more lagrangian particles and years of data as a form of sensitivity analysis, following the recommendations of Brickman and Smith (2002). We determined that the results were representative by calculating the coefficient of variance with the cumulative addition of five years worth of results, where it fell below 5% in all treatments. We then compared the mean and confidence interval of each output to determine the effects of periodicity and depth. We conducted a 2D Kolmogorov-Smirnov (K-S) test (Peacock, 1983; Fasano and Franceschini, 1987) to compare the latitude and longitude distribution of particles released periodically each month and aperiodically. Following the technique outlined by Press et al. (1992), we calculated a K-S statistic ‘D’ that measured the difference between periodic and aperiodic treatments. The statistic is a normalized value between 0 and 1 and was comparable across release, location, depth, and timing. We separately calculated D for each of the 60 monthly release windows over the five years and repeated it at each of the three depths resulting in 180 D values for every site. To detect the presence of an annual signal in D, a discrete Fourier Tranform method (Cooley and Tukey, 1965) was applied to the 60 sequential months of D-values for each site and depth using the numpy package (Harris *et al*., 2020) in Python. From all the sites, we selected several that represent the different geographic areas in the region, for a further comparison of dispersal characteristics and distributions between periodic and aperiodic scenarios.

## Results

The distributional difference between particles released periodicially and those released aperiodically (D) was highly variable depending on the month of periodic release for all sites (Figure 2). There was no consistent pattern of variation in D across years based on the month of release. In most cases there was little to no effect of the dispersal depth on the inter-/intra-annual variability of D on each site. However, those sites contained within the Okinawa Trough had a lower mean D and smaller within-site variability of D at 500m, as did a number of sites in the Northern Izu-Bonin area (Figure 2 b). The most distinct geographical trend in the D values was the on average lower D values in the Okinawa Trough when compared to other regions (Figure 2). At all depths, there is a trend of increasing D values from north to south among the Izu-Bonin sites, while the Mariana sites exhibit an decrease in D values from north to south at a depth of 500m. The distribution of particles aperiodically from each site is consistent with those in close proximity (Figure 3 and 4). For example those particles released from sites in the Southern Mariana area show and East-West distribution while those in the Norther Mariana area have a more typical circular distribution around the source site (Figure 4). Across the Mariana area, constrainement of particles within the vicinity of the source site decreases with depth (Figure 4) while the opposite is true for sites in the Okinawa Tough (Figure 3). The Fourier Transformation detected the presence of an annual frequency in D-values at a number of sites across all geographic areas. However, the strength of these signals did not stand apart from other non-annual frequencies in all cases apart from Nikko Volcano at 100m, Daiichi-Amami knoll at 500m, and South Sarigan Seamount at 1000m. There was no discernable pattern in this annual signal across dispersal depths or source site locations.

**Figure 2.**
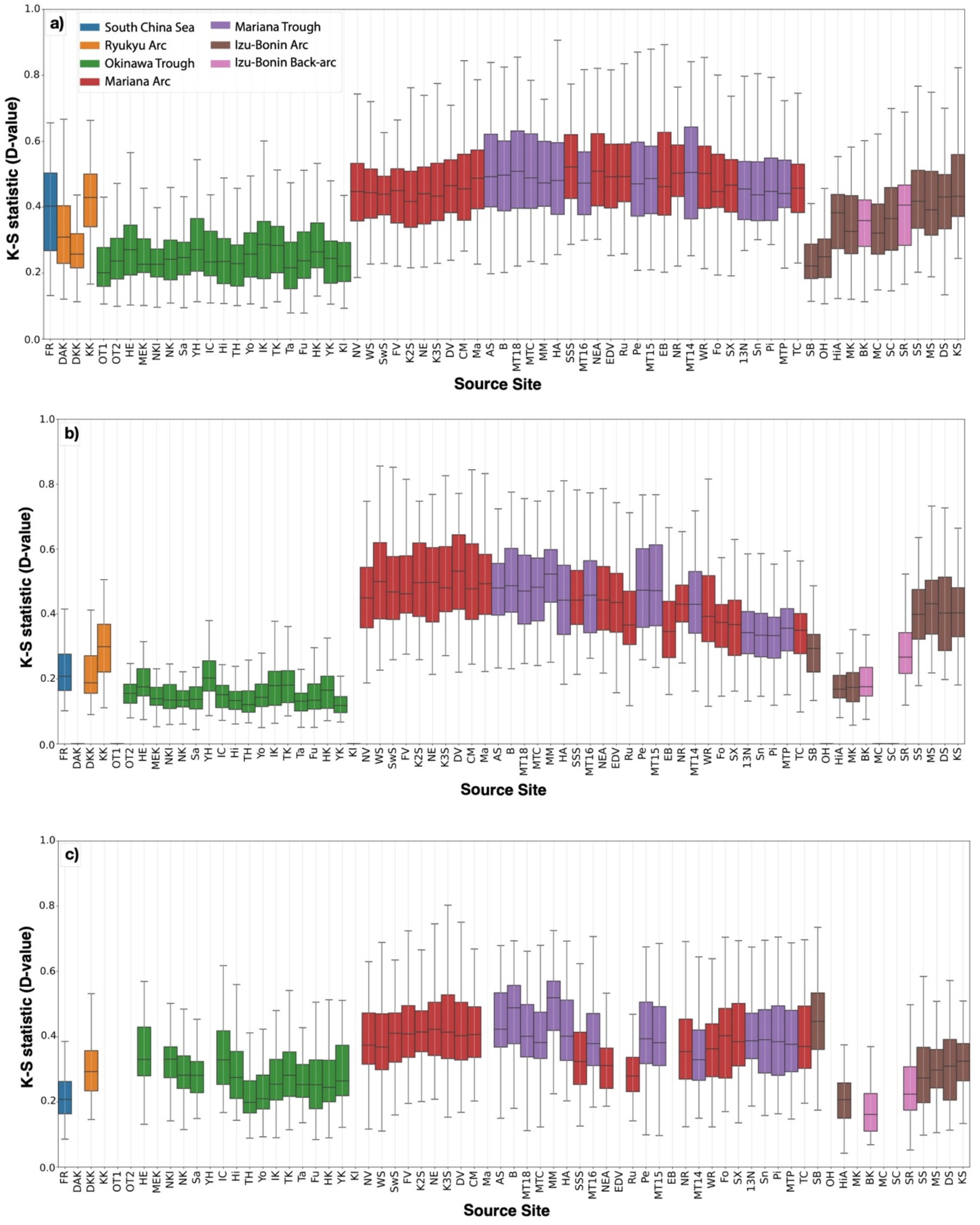
The distribution of the Kolmogorov-Smirnov statistic (D) across all 60 months (5-years) of release for each site. The sites are arranged by their tectonic region and then by lattitude (North – South) to demonstrate the geographical consistency that occurs among depth scenario (a – c). Full names and other metadata for each Source Site in Supplimental material

**Figure 3.**
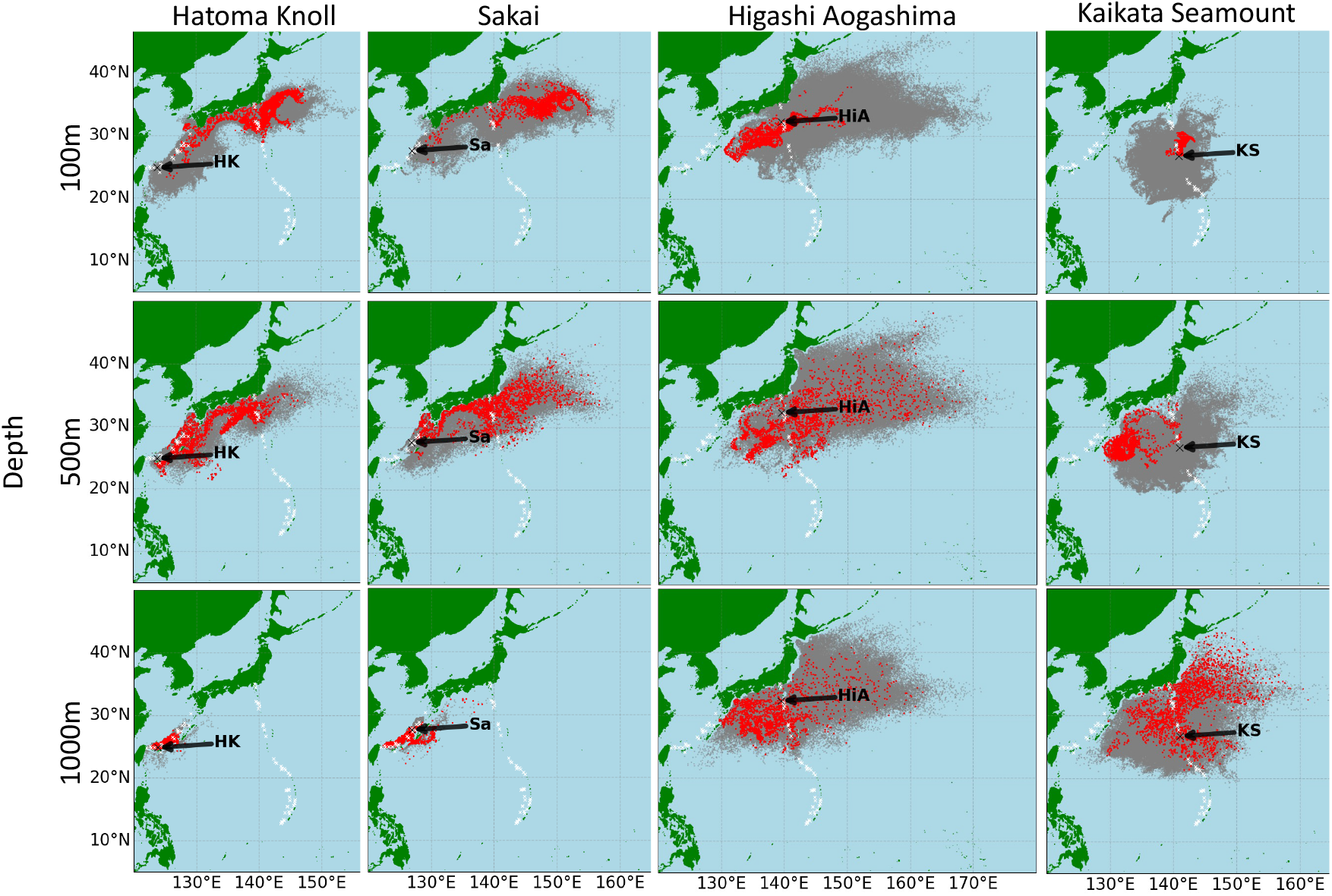
The distribution of Lagrangian particles dispersed aperiodically (grey) and during the single month that resulted in the greatest D-value (red) from the Okinawa Trough or Izu-Bonin area. The source sites (black ‘x’) of the lagrangian particles were selected as representative of the other sites (white ‘x’) within their vicinity.

**Figure 4.**
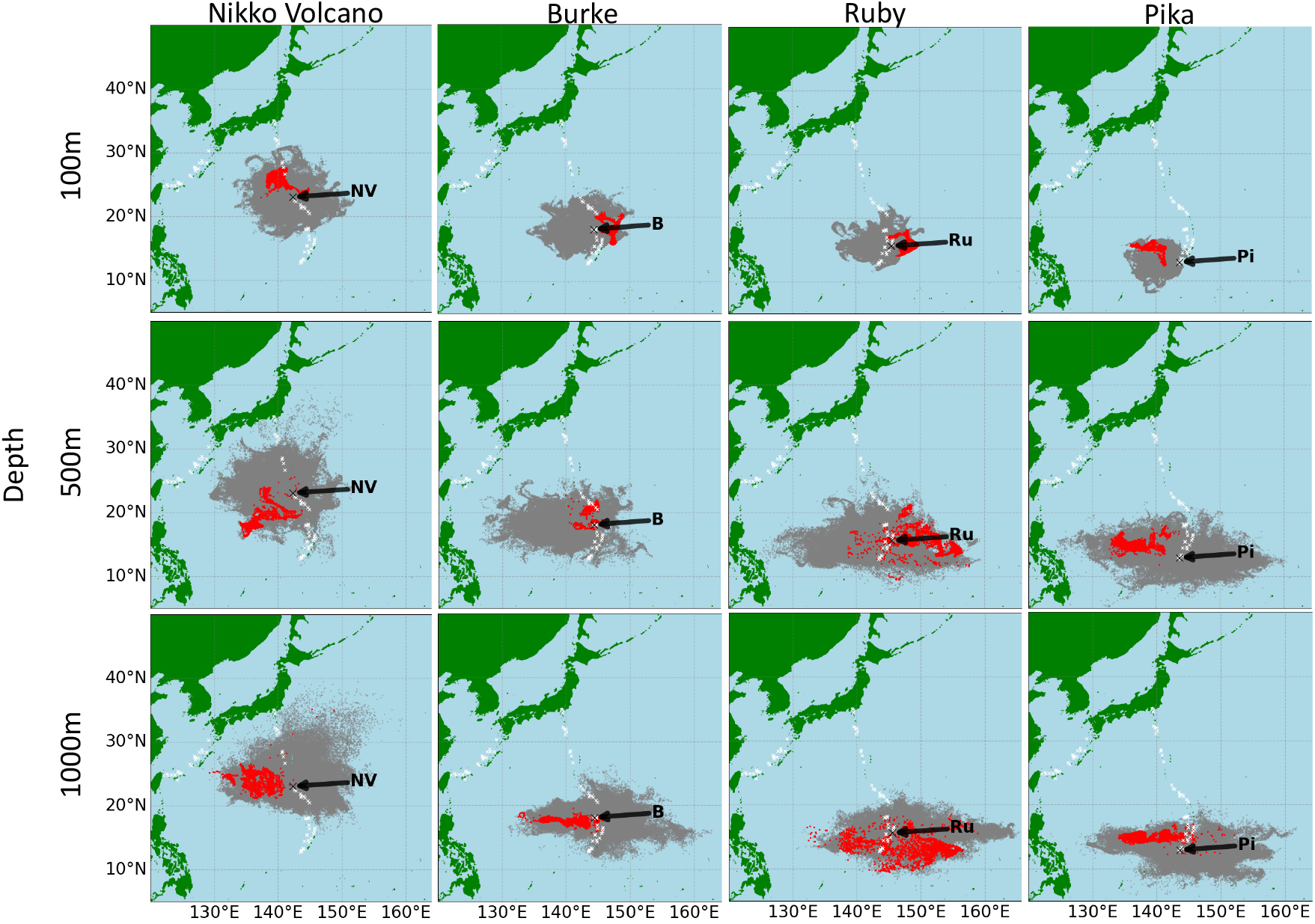
The distribution of Lagrangian particles dispersed aperiodically (grey) and during the single month that resulted in the greatest D-value (red) from the Marianaarea. The source sites (black ‘x’) of the lagrangian particles were selected as representative of the other sites (white ‘x’) within their vicinity.

The dispersal characteristics (PLD, average temperature experienced and total distance travelled) of particles released during the month that resulted in the highest D-value showed no clear or consistent difference with those particles released aperiodically (Figure 5). The monthly variability of PLD, temperature and distance of lagrangian particles was highly dependent on the location of the source site with more adjacent sites showing a similar range (Figure 5). The average temperature, and therefore PLD, was strongly affected by the dispersal depth, much more so than the distance traveled. Lagrangian particles dispersing from all sites aperiodically had an average PLD of 38 at 100 m, 163 days at 500m, and 304 days at 1000m depth. Average temperature experienced decreased with depth as did the variability between sites. The mean total distance travelled by particles did not show a clear relationship with depth and was much more dependent on the location of the source site (Figure 5).

**Figure 5.**
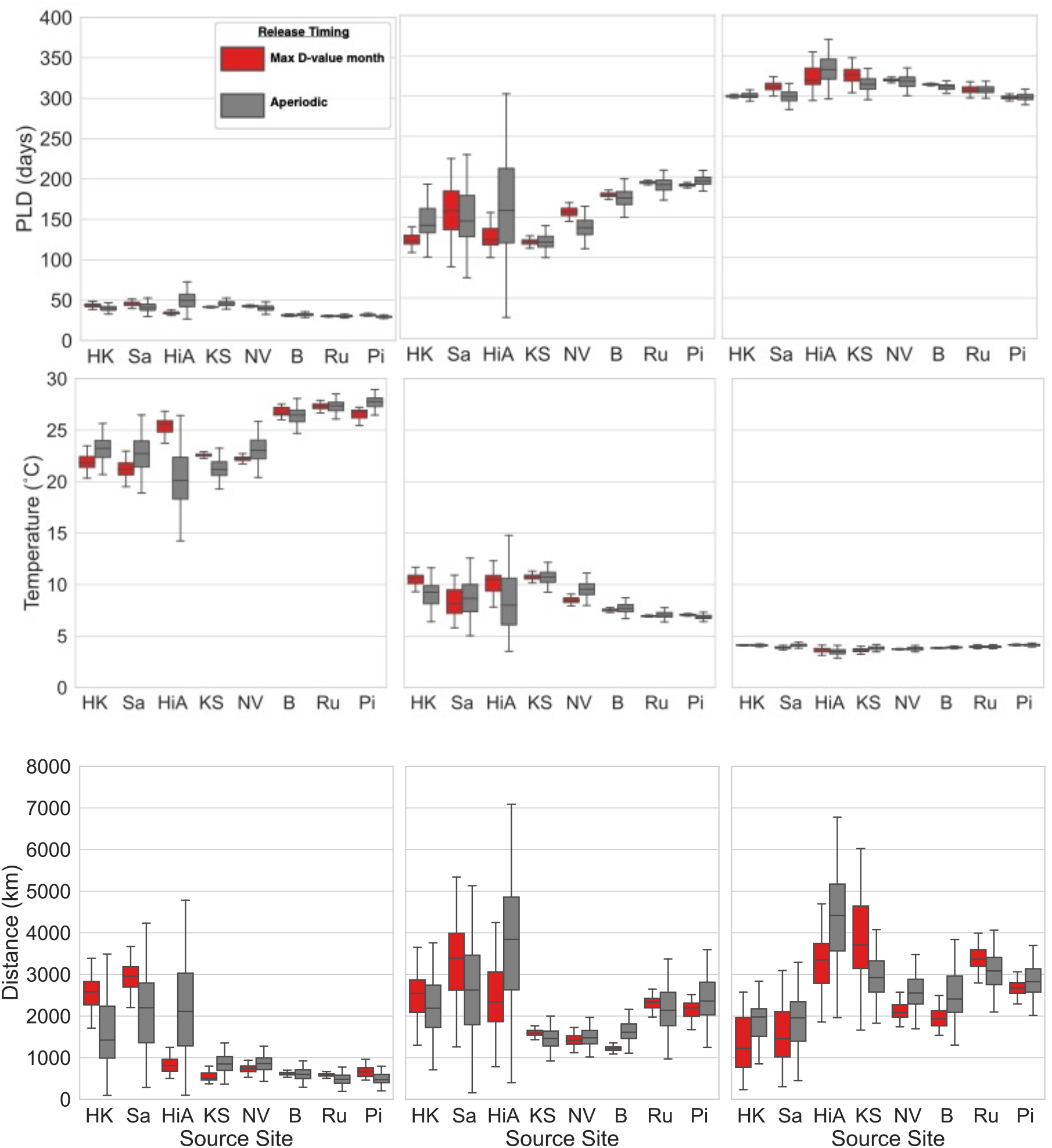
Dispersal characteristics of Lagrangian particles from the example sites shown in figures 3 and 4. There is no consistent difference between the characteristics of particles dispersed from the maximum D-value month (red) and those dispersed aperiodically (grey).

## Discussion

### Dispersal Characteristics

Considering the variability of oceanographic conditions across the region, the effect of periodicity on dispersal distributions and characteristics is highly variable among source sites (Figure 2). A dominant oceanographic feature of this region is the ‘Kuroshio’ western boundary current that travels north within the Okinawa Trough and then Northeast along the eastern coast of Kyushu, Japan. Most particles released from sites in the Okinawa Trough and dispersing at 100 or 500 meters are entrained into the Kuroshio, this is likely the reason for the generally low periodic-aperiodic distributional differences (Figure 2.a and 2.b) as the strong and consistent current constrains their distribution. In the 100 and 500 meter scenarios, the highest D-values for sites in the Okinawa Trough during periodic release months are due to more particles being carried away by the Kuroshio than in the aperiodic scenario, resulting in fewer particles remaining in the Okinawa Trough (Figure 3. a, b, e, f). In the 1000 meter scenarios however, high D-values are caused by a larger than usual proportion of particles leaving the Okinawa Trough through the Kerama Gap (Figure 3. i, j). Particles released from Hatoma Knoll, in the south the Okinawa Trough, are susceptible to being strongly retained in the south at depths of 1000m in certain months (Figure 3. i). The retention of particles in the southern region at 1000m is consistent with the observed distribution of NEMO floats deployed from Hatoma Knoll at the same depth by Mitarai *et al*. (2016).

This retention can even occur at 100m where particles are more often entrained into the Kuroshio and exhibit long-distance dispersal (Figure 3. a). The influence of the Kuroshio can also be seen in the Northern Izu-Bonin sites (Figure3. c, g, k), and in general has a tendancy to decrease the difference in dispersal between periodicly and aperiodicly released particles at 100 and 500 meters. It is perhaps surprising that particles dispersing from vent sites within the path of the Kuroshio experience less variation in dispersal distributions as it has been demonstrated that the Kuroshio exhibits both annual and intra-annual variation in its pathway (Yin *et al*., 2014; Wang *et al*., 2022). The absence of large meandering events from 1990 to 2004 (Qui and Chen, 2021) may result in particularly low or inconsistent inter-annual variation, but this variability is beyond the scope of this study on the effect of intra-annual variability in dispersal based on release periodicity. Our findings suggests that the role of the Kuroshio in driving long-distance transport of lagrangian particles from Okinawa Trough to the Izu-Bonin Arc is consistent regardless of reproductive periodicity at least outside of large meandering years.

In contrast to the effects of the Kuroshio Current, the North Equatorial Current (NEC) appears to have an opposite impact on the affect of periodicity on dispersal distributions. The southernmost sites in the Mariana region, which are most directly affected by the NEC, exhibit a high degree of variability in larval dispersal distribution depending on the timing of release. This trend is observed across all depths tested, but is particularly pronounced in the 500m scenario, where there is a clear reduction in the average D-value further north in the Mariana area, beyond the influence of the NEC (Figure 2. b). In the 100m scenario, the distribution of particles released aperiodically from southern Mariana sites is primarily to the west of the release site (Figure 4. c, d), in contrast to the larger easterly and westerly distribution observed at greater depths (Figure 4. g, h, k, i). Kendall and Poti (2014) previously demonstrated that the seasonality of the NEC contributes to the self-recruitment potential of surface drifters. This seasonality is driven by wind-stress (Qui and Lukas, 1996; Kim et al., 2004), which may explain the greater degree of constraint observed for particles dispersing at shallower depths. The greater spatial distribution of particles travelling at 500 or 1000 meters depth in the Southern Mariana region is likely the result of less frequently reversing current regimes at that depth in combination with the extended planktonic larval duration at the low temperatures experienced. The relative lack of directionality for particles released from the southern Izu-Bonin and northern Mariana sites, as well as the lowed D-values in many cases, is indicative of the absence of prevailing currents in this area. However, the large range in D-values across release months indicates the importance of intra-annual variability on dispersal, even beyond the reach of seasonally fluctuating current systems. The variability beyond the Kuroshio and NEC, and indeed the large variability at all depths, supports the assertations of Adams *et al*. (2011) that surface seasonality can impact the transport of larvae from hydrothermal vents even at great depths.

Based on our results, it is unlikely that temperature variation is the main driver of variations in dispersal distributions, as the temperature range recorded across the extent of this study was very minimal at 1000m (Figure 5). However, it is difficult to separate the role of temperature and ocean currents on dispersal distances because of temperatures influence on PLD in this study. A fixed PLD approach would likely lead to a starker difference in dispersal distance and distributionwith depth (Mitarai *et al*., 2016) and effect the significance of periodicity by not incorporating seasonal temperature variations. Although the resultant PLD of individuals has a large range within this study, all the PLDs of deep-sea species (bar one outlier) collected by Hilario *et al*. (2015) fall within this range.

### Implications for Connectivity

The stochasticity of ocean circulation at depths as great as 1000m, as observed in this study and by driting NEMO floats at this depth (Mitarai *et al*., 2016), results in no annual consistency of dispersal over multi-year scales, as is also the case in surface waters (Siegel et al., 2008). It is unlikely that the addition of more years or even decades of simulatons will result in lower inter-annual variability due to the effects of Kuroshio large meandering events or ENSO (Enfield, 2001). We therefore conclude that at time-scales pertinent to conservation objectives for hydrothermal vents (years – decades), it is not possible to predict the probability of dispersal among hydrothermal vents without sufficient information on species-specific reproductive timing or synchrony. Population genetics studies of vent-associated species in the region, when combined with the results of this study, may provide insights into the role of larval release timing in observed connectivity. For example Xu *et al*. (2018) investigated the genetic connectivity of *Bathymodiolus platifrons* and inferred that the South Okinawa Trough sites of Yonaguni Knoll (YK) and Hatoma Knoll (HK) were source populations for the Mid-Okinawa Trough. Dispersal from the South to the mid Okinawa Trough is consistent across depths but is less likely during months that result in higher entrainement into the Kuroshio at 100m and 500m or in cases where southern retention is strong at 1000m depth. It is certainly possible that this species exhibits periodic spawning as there is evidence for it in species of the same genus (Dixon et al., 2006) as well as other chemosymbiotic mussels (reviewed by Laming *et al*. (2018)) which carry out dispersal in the upper layers of the water column (Arellano et al., 2014). If this species has adapted a periodic reproduction strategy, our findings suggest that this would have minimal impact on its dispersal success as their larvae would consistently be transported north by the Kuroshio if dispersing in the upper layers of the water column, with relatively low intra-annual differences (Figure 2) and no inter-annual consistency. Even in the month that results in the highest deviation from this northern dispersal (low entrainement into the Kuroshio) the particles are mostly lost to the south beyond the vicinity of other suitable habitats or self-recruitment (Figure 3 a). Similarly, the absence of periodicity in the brooding phase of *A. longirostris* from the Okinawa through (Methou et al. in review), in contrast to observations of periodicity in congeneric species from the Atlantic (Copley and Young, 2006; Methou et al., 2022), could be related to the relatively minimal effect of periodicity on dispersal characteristics in this area. It should be noted that no dispersal scenarios tested were able to explain the observations of genetic connectivity from Sagami Bay and the South Okinawa trough to the South China Sea for *B. platifrons* (Xu *et al*., 2018) or *A. longirostris* (Li, 2015; Yahagi et al., 2015). However, certain release months showed a strong and consistent (within the month) Westward directionality in dispersal from the Northern Izu-Bonin area (Figure 3 c and k) and supports the possibility of dispersal in this direction as evidenced from genetic observations of *Gandalfus yunohana* populaitons (Watanabe *et al*., 2020). In this way, observations of intra-annual variability may reveal dispersal processes that can resolve inconsistencies between the directionality of connectivity among hydrothermal vent populations as observed from genetic data and simulations of aperiodic dispersal (e.g. Breusing *et al*. (2021)).

### Conservation

Our results have reveal the effects of release timing on dispersal from hydrothermal vents, which are annually inconsistent and principally constrained by prevailing oceanographic currents or topographic features. Although informed by in-situ and ex-situ observations of vent-associated species, our simulations do not represent a single species or taxonomic group. To accurately represent the dispersal of any hydrothermal vent species through simulations, more research is required into the life history and dispersal behaviour of species, particularly their reproductive periodicity and temperature dependent PLD. Any consequences of periodic reproduction in terms of dispersal potential would be highly dependent on the distribution of the species in question as well as the distribution of suitable habitat. For species that compete for the same local niche space and exhibit similar larval behavior, having differing dispersal periodicity may reduce the chance of regional exclusion through competition and allow them to coexist in the larger metacommunity (Leibold *et al*., 2004; Mullineaux *et* al., 2018; Chase *et al*., 2020). In the Southern Mariana Trough/Arc, larvae are entrained by currents that are highly variable but predominantly disperse them away from other vent sites (Figure 4). In such areas, periodically dispersing species may be particularly vulnerable to low recruitment and more strongly impacted by the effects global climate change has on the seasonality of ocean currents and temperatures in the Pacific (Fukasawa *et al*., 2004). Due to stochastic dispersal on multi-year timescales, predictions of regional impacts from proposed hydrothermal vent mining in the region (Okamoto *et al*., 2019) may not be generalizable across species with different or unknown reproductive periodicity. Therefore, it is recommended to apply the Precautionary Principle (Principle 15, Declaration 1992) and assume the worst-case scenario regarding mining’s disruption of regional dispersal and resultant connectivity.

## Supporting information

Supplemental Table 1 - site metadata

Supplemental Figure 1 - site map

Supplemental Material - Simulation details

