## Supplementary figures and images for "The Role of Reproductive Periodicity in Dispersal Among Hydrothermal Vents and its Implications for Regional Connectivity and Conservation"

### Supplemental Figure 1 - site map

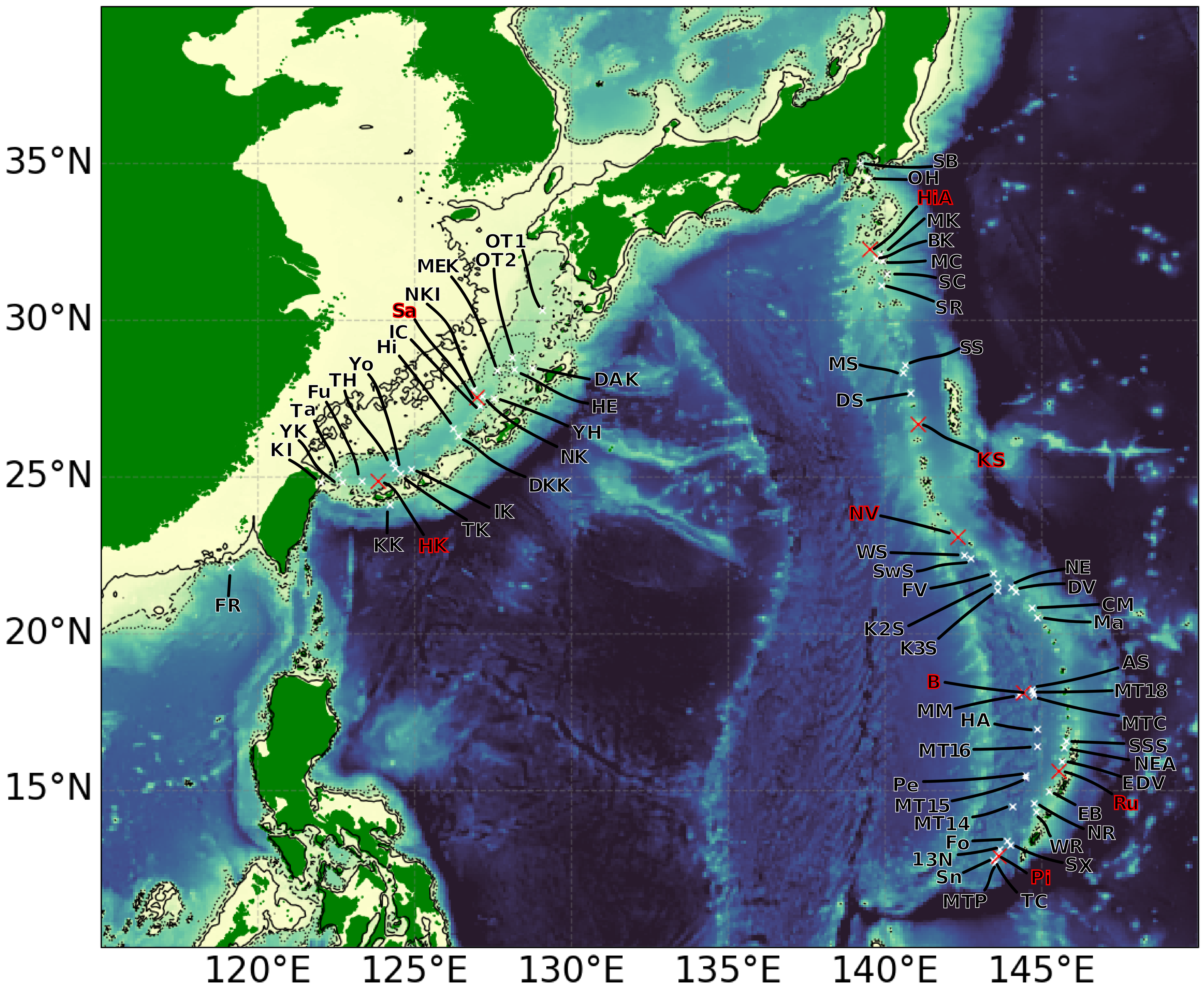
