## Supplemental Material - Simulation details for "The Role of Reproductive Periodicity in Dispersal Among Hydrothermal Vents and its Implications for Regional Connectivity and Conservation"

**Supporting Information: Larval Dispersal Simulation with PARCELS**

An example of the methods described below and all those used in the associated manuscript can be found at:

<https://github.com/otistwo/hydrothermal_thesis/tree/main/PARCELS>

**Ocean Model (Fieldset)**

Three oceanographic variables were used in these simulations (Eastward Ocean Current, Northward Ocean Current and Seawater Temperature) and were freely downloaded from the ‘MOI GLORYS12_FREE’ MERCATOR product. These variables are in the Arakawa C-grid format (ORCA 1/12deg). Additional information on this product can be found at <https://www.mercator-ocean.eu/en/solutions-expertise/accessing-digital-data/product-details/?offer=4217979b-2662-329a-907c-602fdc69c3a3&system=4f9a6cea-7bb0-e4d7-f56d-5dde805195e0>

The horizontal extent of the oceanographic variables and entire simulation was approximately 104°E - 62°W and 0.04°N – 359.95°N, therefore included most of the North Pacific Ocean. The temporal resolution was daily and the extent was five years from 1997/01/01 – 2002/01/01. ‘Ocean PARCELS’ (<https://oceanparcels.org/> ), a series of methods in the python programming language, were used to combine environmental variables into a fieldset, within which simulated particles were released.

**Simulating dispersal with PARCELS**

Simulated particles that represent infinitesimally small parcels of water were released and given behaviors to make them approximate dispersing larvae. Particles were released every six hours from each source site (Supplimental Table 1) for a period and density dictated by the periodicity of the scenario (see methods). Once released into the oceanographic model, particle positions were modified by advection (a 4th order Runge-Kutte implementation) as well as diffusion (Smagorinsky, 1963). The implementation of Smagorinsky diffusion was modified from the tutorial in PARCELS (<https://nbviewer.org/github/OceanParcels/parcels/blob/master/parcels/examples/tutorial_diffusion.ipynb>) to reduce diffusion to 0 at each cell bordering the landmask.

Custom behaviour kernels were attributed to each particle in order to record the distance travelled, average temperature experienced and age at termination. Termination occurred at the end of the Planktonic Larval Duration (PLD), determined by implementing the model of O’Connor *et* al. (2006) as implemented by Mitarai *et al*. (2016). (PLD) = B_0_ − 1.34 × ln (T/Tc) − 0.28 × [ln (T/Tc)]^2^  where T= temperature experienced by the particle at each hourly timestep, Tc was set to 15 and B_0_ = 4.56 as the maximum magnitude of PLD – temperature relationship that fits with the unified model.
